# Identification of senescence-associated drivers of tumour growth and progression using a novel microarray platform

**DOI:** 10.64898/2025.12.20.695739

**Authors:** Hui-Ling Ou, Reuben Hoffmann, Rebecca Smith, Kaylyn Devlin, Mark Dane, Daniel Muñoz-Espín, James E Korkola

**Author notes:** H-L.O. and R.H. contributed equally to this article. D.M-E. and J.E.K are co-senior. Author’s Contributions: H-L.O and R.H. designed and performed experiments, performed analysis, and contributed to the writing of the manuscript. R.S. and K.D. produced the MEMA and assisted in the MEMA experiments. M.D. performed analysis of the MEMA data. D.M.E. and J.E.K. conceived of the study, designed experiments, wrote the manuscript, and provided funding support for the project. J.E.K. also performed experiments and data analysis.

## Abstract

Senescence and the senescence associated secretory phenotype (SASP) are implicated in promoting early tumorigenesis but due to the complexity of SASP it has been difficult to identify the responsible factors. We used canonical SASP factors on our microenvironment microarray (MEMA) platform to systematically identify SASP-associated drivers of tumorigenesis in breast and lung cancer cells. We found multiple SASP factors enhanced the proliferation and overall cell numbers for both lung and breast cells grown on the MEMA, and that there was significant overlap in SASP-associated growth-promoting factors between the two different cell types. We validated the ability of several factors, including IL-6, TGF-β and EGF, to drive growth in *in vitro* assays. Interestingly, these factors were effective in driving growth and survival in cells that were altered (either immortalized or fully transformed) but not in normal cells and impacted breast cells differently depending on the age of the patient. RNAseq identified upregulation of wound-healing and stem-cell programs in SASP factor-treated cells. Many of these same SASP factors were present in conditioned media collected from senescent cells, which enhanced the growth of both lung and breast cancer cells, and inhibitors of the specific SASP factors partially reduced growth. Similarly, targeted inhibition of EGF partially reduced lung tumour growth in xenografts when senescent but not normal fibroblasts were co-implanted. Our findings have identified core SASP drivers of tumorigenesis and suggest that effective tumorigenesis driven by SASP is multifactorial and requires alterations in the target cells to achieve maximal response.

## Introduction

Multiple lines of evidence suggest the local tissue microenvironment plays a crucial role in early tumorigenesis. A healthy microenvironment has been proposed to suppress the growth of pre-neoplastic cells, while a pro-tumorigenic microenvironment can promote the growth and progression of those same cells (Laconi *et al*, 2020). A number of different conditions have been implicated in creating a pro-tumorigenic microenvironment, including chronic inflammatory conditions. For example, pancreatitis and colitis are both associated with a significantly increased risk of cancer development (Chung *et al*, 2012; Lakatos & Lakatos, 2008). However, the greatest risk factor for cancer formation is aging (Marongiu & DeGregori, 2022). While aging can lead to an increased mutational burden, it is also associated with tissue dysfunction. In particular, the number of senescent cells present in tissues significantly increases with aging (Omori *et al*, 2020; Ovadya *et al*, 2018; Pignolo *et al*, 2020).

Senescence is a form of durable cellular arrest that can be triggered by a variety of stimuli, including telomere loss, oncogene activation, cellular stresses, and external cellular signalling (Campisi & d’Adda di Fagagna, 2007; Ou *et al*, 2020). Senescent cells differ in several ways from normally-growing or quiescent cells, one of the most notable of which is the implementation of a potent Senescence-Associated Secretory Phenotype (SASP). SASP results in increased expression and secretion of various cellular signals like interleukins, chemokines, growth factors, and matrix metalloproteinases and tissue remodelling factors; these signals have strong ties to inflammation, immune cell infiltration and modulation, growth regulation, and microenvironment remodelling (Campisi & d’Adda di Fagagna, 2007; Ou *et al*., 2020).

Senescence has long been referred to as a “double-edged sword” or antagonistic process (Campisi, 1997; Liu *et al*, 2018; Ohtani *et al*, 2012). In cancer, it is considered a tumour suppression mechanism, as damaged cells that become senescent are prevented from propagating and becoming cancerous, and through SASP signalling cause surrounding cells that may also be damaged or undergo transformation to become senescent (Campisi, 1997; Liu *et al*., 2018; Ohtani *et al*., 2012; Ou *et al*., 2020). Indeed, to become cancerous, a cell must either avoid or escape from senescence. In normal tissue, senescent cells exist transiently and are cleared by immune cells (Kale *et al*, 2020; Ovadya *et al*., 2018). However, a large pool of evidence shows a detrimental, pro-tumorigenic effect of the chronic persistence of senescent cells that can arise through a variety of means, including aging and response to chemotherapy (Demaria *et al*, 2017; Ou *et al*., 2020). High senescent cell count in tumours is associated in some cases with a worse prognosis (Pare *et al*, 2019) and, experimentally, senescent cells can drive tumorigenesis (Krtolica *et al*, 2001) and alter therapeutic response (Demaria *et al*., 2017). It is hypothesized that these effects are, at least in part, driven by changes in the local microenvironment via SASP signalling (Campisi & d’Adda di Fagagna, 2007; Collado & Serrano, 2010; Coppe *et al*, 2010; Ou *et al*., 2020).

Although it is well-accepted that senescent cells and chronic SASP likely support tumorigenesis, it has been difficult to define the specific SASP factors which are most important for driving the growth and progression of pre-neoplastic cells. This is due to both the complexity and heterogeneity of the SASP, which produces a myriad and dynamic range of secreted factors that can vary significantly between tissues (Hernandez-Segura *et al*, 2017; Ou *et al*., 2020). In an attempt to more systematically address the role of the SASP, we utilized our novel microenvironment microarray (MEMA) platform (Smith *et al*, 2019; Watson *et al*, 2018) to interrogate a library of common SASP factors on pre-neoplastic and neoplastic cell growth. As models, we utilized lung cancer cells (A549) and human mammary epithelial cell (HMEC) cultures that were isolated from normal patients undergoing reduction mammoplasties. The HMEC cultures have been modified with sequential addition of oncogenic constructs to give rise to a continuum of cells ranging from primary to post-stasis to immortalized but not fully transformed cells.

We report here our results from the MEMA platform analysis of interactions between combinations of extracellular matrix factors and secreted ligands, which allowed us to identify prospective —context dependant—“hits” among the SASP factors that contributed to cell growth and survival. We validated the growth and survival effects of these hits in our cell lines of interest, revealing both similarities and distinctive features between the lung and breast samples. Our findings demonstrate tissue of origin effects, effects on growth related to the degree of transformation, and effects related to the age of the HMEC donor all playing a role in determining how SASP may drive the growth of early tumour cells. As a functional validation and *proof-of-concept*, we further demonstrate the presence of these same SASP factors in conditioned medium from cells induced to become senescent by irradiation or chemotherapy treatment, and the ability of targeted inhibitors against these SASP factors to block the growth of recipient cells both *in vitro* and *in vivo*. Of note, RNAseq analysis identified induction of key tumorigenesis-associated programs, including wound healing and stem cell induction. Our use of the novel MEMA technology identifies key SASP factors that are likely to contribute to a pro-tumorigenic microenvironment to drive the growth and progression of early cancerous cells and the potential to target these to intercept early tumorigenesis.

## Materials & Methods

### Cell lines and reagents

The human lung adenocarcinoma cell A549 was obtained from American Type Culture Collection (ATCC) and cultured in DMEM growth medium supplemented with 10% Foetal Bovine Serum (FBS). Human Mammary Epithelial Cell (HMEC) primary cells and derived cell lines were obtained from Martha Stampfer of Lawrence Berkeley National Laboratory and cultured in complete M87A media as described previously (Garbe *et al*, 2014; Romanov *et al*, 2001); a complete list of HMEC lines used is provided in **Supplemental Table 1**. Human Pulmonary Fibroblasts from adults (HPF-a) were purchased from ScienCell and cultured in fibroblast medium (FM, #2301, ScienCell) supplemented with 2% FBS, 1% fibroblast growth supplement (#2352, ScienCell), 200 units/ml of Penicillin and 200 μg/ml of Streptomycin (#0503, ScienCell). Human bronchial epithelial cells were obtained from ScienCell and cultured in bronchial epithelial cell medium (BEpiCM, #3211, ScienCell) supplemented with 1% bronchial epithelial cell growth supplement (#3262, ScienCell), 200 units/ml of Penicillin and 200 μg/ml of Streptomycin (#0503, ScienCell). For tumour imaging of *in vivo* experiments, A549-Luc2 cells was obtained from ATCC and cultured using the same condition as A549 cells. All cells were maintained at 37°C incubators with 5% CO_2_ and under regular monitoring for Mycoplasma contamination.

### Cell Line Treatments

Cellular senescence was induced by treating HPF-a with 2.5 μM cisplatin (CDDP, Sigma-Aldrich) for 10 days or with 10-Gy ionising radiation (195 kV, 10 mA, Xstrahl RS225M X-ray Generator), and in HMEC by treating with 10 nM paclitaxel (Selleckchem), 50 nM doxorubicin (Selleckchem), or 5 μM cisplatin (Sigma-Aldrich). Inhibition of epidermal growth factor (EGF)-promoted cellular growth was accomplished by treating A549 cells with gefitinib (ZD-1839, Iressa, Selleckchem) at the indicated concentrations and TGF-β promoted growth was inhibited by treating HMEC cells with galunisertib (Selleckchem) at the indicated concentrations.

### Colony formation assay

HMEC cells were seeded at a density of 500 cells/well and A549 cells were seeded at a density of 750 cell/well in 6-well plates and allowed to grow for the indicated times. Wells were washed once with PBS prior to fixation with 4% paraformaldehyde (#43368, Thermo Fisher Scientific) for 5 minutes. Fixed cells were washed twice with PBS and immersed in cold methanol for permeabilisation for 10 minutes. The cells were then left to dry for 20 minutes and subsequently stained with 0.5% crystal violet staining solution (0.5 g crystal violet powder, Sigma-Aldrich, dissolved in 20 % methanol) for 30 minutes. Once staining was completed, cells were washed 3 times with water to eliminate any residual stain and left to dry before imaging. The colony counting was performed utilising Clono-Counter software (Niyazi *et al*, 2007) by considering both the size and density per colony. In addition to colony counting, the stain was eluted with 10% acetic acid and subjected to optical density (OD) measurement at the wavelength of 595 nm with Infinite 200 PRO Multimode Spectrophotometer (PECAN) or GloMAX (Promega) microplate readers. The readings were then used for calculating the clonogenic potential per treatment by normalising with corresponding control.

### Conditioned media collection & treatment

Ten days after the induction of senescence (see cell line treatments), growth media was removed and cells were washed twice with PBS to eliminate any residual conditioning or chemotherapeutic. Low-serum (0.4% FBS) media was then added to cells for 24, 48- or 72-hour incubation before collection. The harvested conditioned media were centrifuged at 500 x g for 3 min to precipitate any cells or cell debris before being used for further treatment.

### Microenvironment microarray (MEMA)

MEMA plates were prepared as described previously (Smith *et al*., 2019; Watson *et al*., 2018) by printing spots of ECM factors at varying concentrations into each well of an 8-well plate (Nunc) using an Arrayer Robot (Aushon). The full list of factors utilized for these experiments is available as **Supplemental Table S1**. MEMA assays were performed by plating A549 or 153D1MY (HMEC, post-menopausal donor, immortalized) cells in cell-line-specific low-serum media, allowing them to adhere overnight, and then spiking in SASP factors at varying concentrations (or PBS control; see Table [O-S2]), and allowing 72 hours of growth. At the end point, EdU was spiked in one hour prior to fixation in 4% PFA for 15 min. Fixed plates were permeabilized in Triton-X100 (Sigma) and stained with Click-IT EdU (Life Technologies), DAPI, and selected antibodies as indicated in the text. Stained plates were imaged using a HCA microscope system (Nikon) and analysed using custom R scripts (see below).

### MEMA Analysis

MEMA analyses were performed as described previously (Smith *et al*., 2019; Watson *et al*., 2018). Briefly, we pre-processed and normalized data using open-source R software (see similar pipeline at https://github.com/HeiserLab/TNBC_MEMA). The cell count in each MEMA spot was based on the DAPI stained nuclei. EdU intensity was auto-gated to label cells as EdU^+^ and the proportion of EdU^+^ cells in each spot was reported to measure proliferation. The per-cell intensity values were median summarized to the spot level. Each intensity and morphology signal was independently RUV normalized in a series of matrices with arrays as the rows and spots as the columns (Hunt *et al*, 2020, 2022). The RUV controls were the residuals created by subtracting the replicate median from each spot value. After RUV normalization, bivariate LOESS normalization was applied to the normalized residuals using the array row and array column as the independent variables. After normalization, the ∼15 replicates of each condition were median summarized to the MEP level. Major findings from the MEMA were recapitulated in at least 3 experimental replicates. Dunnette’s test was run with the DunnettTest function from the R package DescTools using the PBS spots as the controls to identify significant microenvironment factors.

### Live-cell imaging and analysis (Incucyte)

Cells were inoculated onto plates coated with or without collagen I (Cultrex) at 200 μg/ml and left overnight before treatment. Real-time cell growth was monitored using IncuCyte® SX5 Live-Cell Analysis Instrument (Sartorius AG). For A549, cells were seeded at a density of 25,000 cells per well in a 24-well plate the day before treated with designated SASP ligands, and growth was monitored for 3-7 days. For the HMEC cells, 10,000 cells were seeded per well in a 96 well plate, then treated 24h later with the designated ligands for 3-7 days. Cell confluency was analysed using the IncuCyte® software and calculated as percentage over the baseline confluence.

### Proteomic analysis of conditioned media

Conditioned media (CM) from control or senescent cells were harvested after 72-h incubation with low-serum growing media. CM were centrifuged at 500 x g for 3 minutes to eliminate any residual cell debris before being subjected to Proteome Profiler Human XL Cytokine Array kit (#ARY022B, R&D Systems). Arrays were visualised with ECL Prime Western Blotting detection reagents (#RPN2232, Cytiva) for the chemiluminescent signals. Images were captured using ChemiDoc™ Imaging System (Bio-Rad). For evaluating specific secretomes, ELISA kits were used for determining the amounts of EGF1(#DEG00, R&D Systems) or TGF-β1 (#DB100C, R&D Systems) in indicated CM with normalization based on cell content.

### RNA extraction (for RT-qPCR & sequencing)

RNA extraction of cells was performed using either Trizol (life Technologies) or the Monarch® Total RNA Miniprep Kit (#T2010, New England BioLabs) following manufacturer’s protocols. For the kit, cells were washed once with PBS before RNA lysis buffer was added to the cells. Cells were then scraped off the dish and subjected to genomic DNA removal with the gDNA removal column provided in the kit. The flow-through containing RNA was treated with equal volume of ethanol (≥ 95%) for precipitation and loaded to the RNA purification column. The RNA-containing column was washed once with RNA Priming Buffer followed by two-time washing with RNA Washing Buffer. After eliminating thoroughly the Washing Buffer, 40 μl nuclease-free water was added to the column for RNA elution. The quality and quantity of extracted RNA was determined using NanoDrop One (Thermo Scientific).

### RNA-sequencing

RNA with indicated treatment were sent to BGI Hong Kong for performing RNA sequencing following the strand-specific transcriptome library construction protocol (DNBSEQ G400 platform) with read length of PE100. In brief, mRNA was purified from total RNA using oligo(dT)-attached magnetic beads and subjected to fragmentation, from which first- and second-strand cDNA were synthesised, followed by end repair and adaptor ligation. The amplified products from further PCR reaction were validated on the Agilent Technologies 2100 bioanalyser prior to circularisation and sequencing. BGI performed initial sequence alignment and normalization by FPKM (Fragments Per Kilobase per Million mapped fragments) methodology. The FPKM values were log transformed and median centered prior to clustering, differential gene expression, and Gene Set Enrichment Analysis (GSEA), which were performed as described previously (Smith *et al*, 2020). Briefly, clustering and visualization of the most variable genes (highest standard deviation across all samples) was performed using standard approaches (Eisen *et al*, 1998). R modules were used for differential gene expression using mult-test and the Benjamini-Hochberg method to correct for multiple comparisons (Pollard KS, 2010) and GSEA using ClusterProfiler (Yu *et al*, 2012) utilizing GO and KEGG databases to define genesets.

### RT-qPCR

Total RNA extracted from cells was first used as templates for cDNA synthesis with High-Capacity RNA-to-cDNA™ Kit (#4387406, Applied Biosystems). For each reverse transcription (RT) reaction, 2 μg RNA was mixed with RT enzymes and the corresponding buffer before loaded into the thermal cycler, which was programmed as 60-min incubation at 37°C followed by 5-min incubation at 95°C. The cDNA products were then diluted 4 times with nuclease-free water and stored at -20°C or directly used for quantitative PCR.

To prepare for quantitative PCR, 9 μl cDNA (< 100 ng in total) was mixed with Luna^®^ Universal qPCR Master Mix (#M3003, New England BioLabs) and corresponding primer set (final concentration 0.25 μM) before dispensed into a 96-well. The sequence of primers used in this study are: *ACTB* fw-AGAAGGATTCCTATGTGGGC, rev-TACTTCAGGGTGAGGATGC; *CCL20* fw-GCTGTACCAAGAGTTTGCTC, rev-AGTTGCTTGCTTCTGATTCG; *CXCL1* fw-CCCAAGAACATCCAAAGTGTG, rev-CATTCTTGAGTGTGGCTATGAC ; *CXCL8* fw-ACTCCAAACCTTTCCACCC, rev-CAATAATTTCTGTGTTGGCGC;*CXCL10* fw-ACGTGTTGAGATCATTGCT, rev-AAATTCTTGATGGCCTTCGA; *CDKN1A* fw-TCTTGTACCCTTGTGCCTC, rev-GGTAGAAATCTGTCATGCTGG; *CDKN2A* fw-GAAGGTCCCTCAGACATCC, rev-GTAGGACCTTCGGTGACTG; *GAPDH* fw-TCAAGATCATCAGCAATGCC, rev-CGATACCAAAGTTGTCATGGA; *IL1A* fw-GACGGTTGAGTTTAAGCCA, rev-GCTTGATGATTTCTTCCTCTG; *IL6* fw-GATTCAATGAGGAGACTTGCC, rev-TGTTCTGGAGGTACTCTAGGT; *LMNB1* fw-GTATGAAGAGGAGATTAACGAGAC, rev-TACTCAATTTGACGCCCAG; *MMP3* fw-GACTCCACTCACATTCTCC, rev-AAGTCTCCATGTTCTCTAACTG; *MMP9* fw-TGCAACGTGAACATCTTCG, rev-GAATCGCCAGTACTTCCCA; *TP53* fw-CTCAGATAGCGATGGTCTGG, rev-CTGTCATCCAAATACTCCACAC; The reaction plate was then loaded to QuantStudio™ 1 Real-Time PCR System (#A40427, ThermoFisher Scientific), which was programmed to SYBR^®^ scan mode with the following settings: initial denaturation at 95°C for 60 seconds (x1 cycle) followed by 40 cycles of denaturation (95°C for 15 seconds) and extension (60°C for 30 seconds) with plate reading per cycle, and ultimately the melting curve between 60-95°C (x1 cycle).

### Senescence-associated β-galactosidase assay (SA-β-Gal assay)

Activity of β-galactosidase was detected using Senescence β-Galactosidase Staining Kit (#9860, Cell Signaling) as described in the protocol. In brief, cells seeded in 6-well plates were washed once with PBS before fixed with 1X Fixative Solution for 10-15 minutes. Meanwhile, the β-Galactosidase Staining Solution was prepared and pH adjusted following the instruction of protocol. After fixation, cells were washed twice before subjected to staining at 37°C. Depending on cell types and senescence inducers, the incubation time for staining ranged between 8-16 h. The staining solution was then replaced with PBS and the plates were imaged with Zeiss Axio Observer 7 microscope and ZEN Blue software (ver 3.1). Positive cells for SA-β-Gal was determined by manually assessing randomised 6-9 images per group with on average more than a thousand cells counted for each experiment.

### Western blot

Cells were harvested using protein lysis buffer consisting of RIPA buffer (#R0278, Sigma-Aldrich), proteinase inhibitor cocktail (#04 693 132 001, Roche), phosphatase inhibitor cocktail (#04 906 837 001, Roche) and 1mM EDTA. Cells were collected with cell scrapers and incubated at 4°C with agitation (1400 rpm) for 30 minutes. Cell Lysates were then centrifuged at 12500 x g to precipitate any cell debris. The clear supernatant was then transferred to a new tube and stored at -20°C or used directly for quantifying the protein concentration with Pierce™ BCA Protein Assay kit (#23225, Thermo Scientific). A total of 30 μg protein was then mixed with NuPAGE™ LDS sample buffer (#NP0008, Invitrogen) and boiled at 95°C for 15 minutes before loaded on the precast Mini-PROTEAN TGX gels (#4561083, Bio-Rad). Separation of proteins was achieved by electrophoresis at 120 V for 1 hour in Tris/Glycine/SDS buffer (#1610772, Bio-Rad) while the transferring of protein to PVDF membranes (#IPFL00005, pore size 0.45 μm, Merck Millipore) was carried out using settings of 15 V for 16 hours at 4°C in Tris/Glycine buffer (#1610734, Bio-Rad) with 20% methanol. The membrane was then blocked using 5 % skimmed milk dissolved in TBS for 1 h at room temperature. The primary antibodies used in this study are as following: phospho-Rb (S807/811, #9308, Cell Signaling); phospho-p53 (S15, #9284, Cell Signaling; S20, #9287, Cell Signaling); p53 (DO-1, #sc-126, Santa Cruz Biotechnologies; DO-7, #sc-47698, Santa Cruz Biotechnologies); p21 (#556431, BD Pharmingen) and GAPDH (#10494-1-AP, Proteintech). Peroxidase-conjugated secondary antibodies (anti-mouse, #715-035-151; anti-rabbit, #711-035-152, Jackson ImmunoResearch) were used in combination with ECL Prime Western Blotting detection reagents (#RPN2232, Cytiva) for detecting chemiluminescent signals. Images were captured using ChemiDoc™ Imaging System (Bio-Rad) with optimised exposure times and analysed with ImageJ (ver1.53s, National Institute of Health, US).

### Cell viability assays (IC50 and GR50)

The half-maximal inhibitory concentration (IC50) of the EGFR blocker gefitinib on A549 cells was determined by testing cell survival over a range of gefitinib dilution. A549 cells were seeded at a density of 6000 cell per well in a 96-well plate the day before treated with series dilution of gefitinib. After 72-h treatment, CellTiter-Blue^®^ (#8080, Promega) was added in a 1:40 dilution for detecting the cell viability. After 2 h incubation at 37°C, the fluorescence signal was measured at excitation 560 nm/emission 590 nm with Infinite M200 Pro plate reader (TECAN Trading AG). For assessment of galunisertib response in HMEC cells, we used CellTiterGlo (CTG, Promega) as described previously (Heiser *et al*, 2012). Briefly, 10,000 HMEC cells were plated into 96-well plates and drug was added the next day at six different concentrations were applied in triplicate. After 72 h, CTG was added and luminescence was read on a GloMax (Promega) instrument, comparing to time 0 controls. Growth rates (GR) and the concentration required to inhibit growth by 50% (GR50) were calculated as previously described (Hafner *et al*, 2016).

### Xenograft experiment

For xenograft experiments, suspension of A549-Luc2 cells and HPF-a cells were prepared in PBS and mixed in 1:1 with Matrigel^®^ Matrix (#356234, Corning). A total of 4 million A549-Luc2 cells were injected subcutaneously in A549-only control group while a combination of 4 million A549-Luc2 cells and 1 million of CDDP-induced senescent HPF-a or normal HPF-a cells were co-injected in the corresponding A549+HPF-a groups. The growth of xenografts was monitored twice per week with digital calliper once they were palpable. Tumour volume was calculated using the formula X^2^*Y/2 where X is the width and Y is the length of tumour. Treating regiment of animals with Gefitinib (ZD1839) were initiated 2 weeks after xenograft transplantation, in which Gefitinib (50 mg/kg body weight) or vehicle was administrated via oral gavage 5 days per week for 3 weeks consecutively. In addition to volume, tumour growth derived from injected A549-Luc2 cells was monitored using IVIS Spectrum Xenogen imaging system (Caliper Life Sciences) and analysed with Living Image^®^ software (ver 4.5.5), for which animals were injected intraperitoneally with D-Luciferin (150 mg/kg body weight, #122799, Perkin Elmer) 10-15 minutes prior to imaging. The tumour growth trend was assessed by normalising the luminescence radiance per timepoint to the baseline signal whilst the growth ratio (%) was generated by normalising the final radiance increased to the baseline signal.

Animals were sacrificed at the end of experiment following the approved study plans or when reaching the humane endpoint where the average diameter of tumour [(X+Y)/2] exceeded 1.5 cm. All animal works in this study were approved for Ethical Conduct by the Home Office England and Central Biomedical Services under the regulation of the Animals Act 1986 complying with The International Guiding Principles for Biomedical Research Involving Animals.

### Statistical Analyses

Information on biological replicates (indicated as N) and technical replicates can be found in the respective figure legends. The reported statistics used sample means, standard deviation (SD), standard error of the mean (SEM), and p-values obtained from unpaired parametric *t*-tests of sample sizes of equivalent variance (unless otherwise noted in figure legends) or ANOVA analysis.

## Results

### MEMA experiments

SASP has long been implicated in driving pre-neoplastic cell growth and progression to invasive disease (**Fig.1a**), but the factors involved remain incompletely understood (González-Gualda *et al*, 2022; Haston *et al*, 2023). We initiated our studies by examining the growth of A549 and the HMEC cell strain 153LD1MY on our MEMA platform. MEMA allow for the comparison of thousands of unique combinations of extra-cellular matrix proteins and soluble ligands for their effects on cellular phenotypes. We selected ligands that have previously been implicated in senescence for testing on the platform, resulting in 63 ligands and 25 ECM proteins for a total of 1,575 distinct combinatorial conditions that were interrogated for their impact on cellular growth. We allowed the A549 and 153LD1MY cells to adhere overnight on the MEMA platform. For A549 cells, we replaced the medium the next day with low serum (0.1% FBS) containing medium to better mimic *in vivo* conditions and then added ligands. For 153LD1MY cells, we replaced the medium the next day with fresh full medium, as these non-transformed epithelial cells do not grow as robustly as A549 cancer cells, and then added ligands individually to each well. The 153LD1MY and A549 (low serum) cells were left for 72h on the MEMA prior to fixation for immunofluorescent (IF) analysis (EdU was added to the medium 1h prior to fixation). Our primary endpoints were cell counts (as measured by image segmentation of DAPI stained nuclei) and EdU incorporation for measurements of cell proliferation.

**Figure 1.**
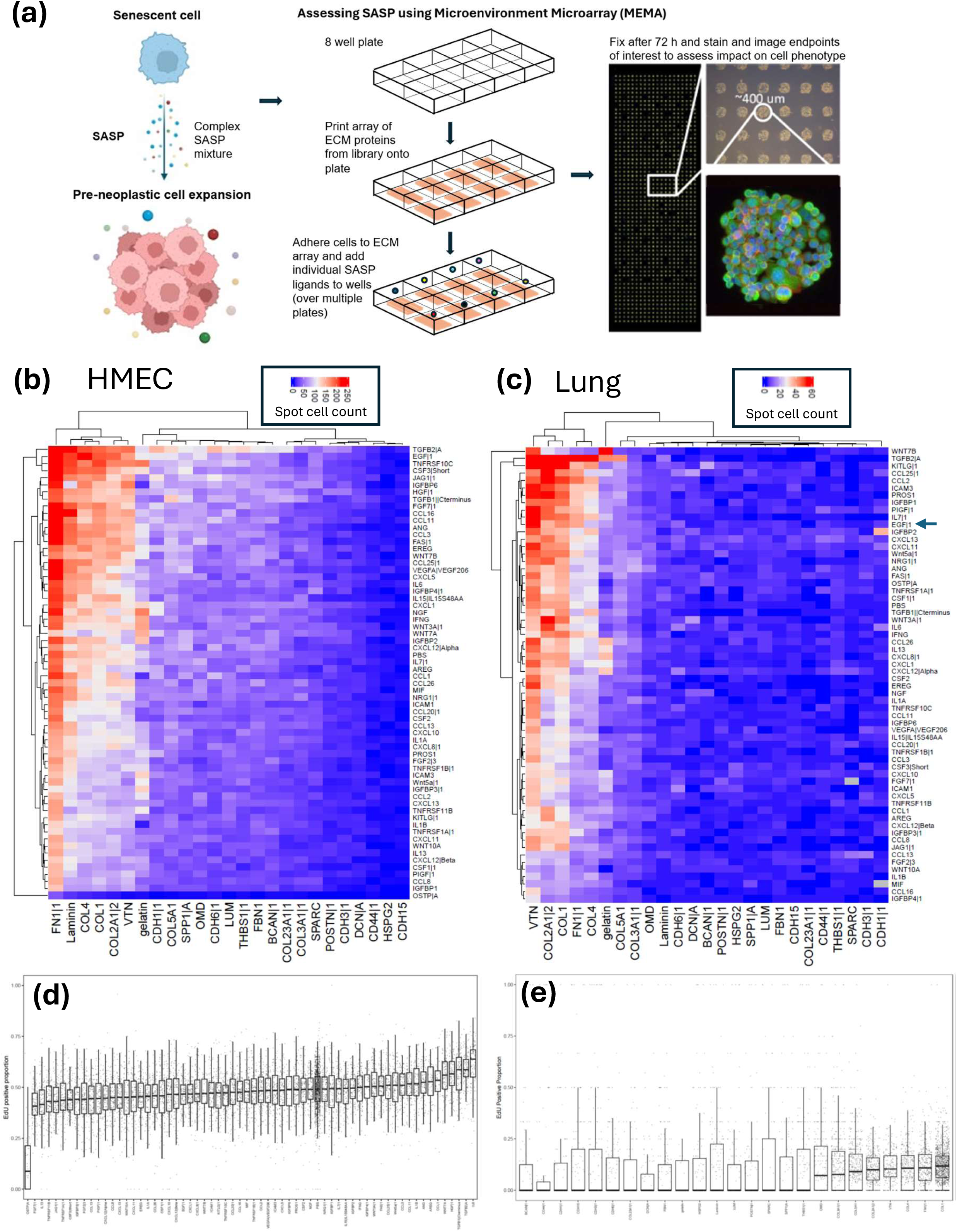
Microenvironment microarray (MEMA) facilitates identification of senescence secretory phenotype factors (SASPs) involved in promoting proliferation of human mammary epithelial cells (HMECs) and alveolar lung adenocarcinoma A549 cells. **(a)** SASP as a driver of pre-neoplastic cell growth and progression. Following unrepairable stress or damage, cells can become senescent and drive expansion of preneoplastic cells through SASP. The exact SASP factors that are most responsible for driving this growth remain incompletely understood. **(b)** Spot cell count for HMEC cells showing conditions that enhance cell numbers **(c)** Spot cell count for A549 cells showing conditions that enhance cell numbers. **(d)** EdU positive proportion, ranked by ligand from low to high, for the HMEC cells. **(e)** EdU positive proportion, ranked by ECM from low to high, for the A549 cells.

We prepared heatmaps of cell number as a function of the ECM and ligand for each condition interrogated (**Fig.1b, 1c**) for both A549 and 153LD1MY cells. This initial view of the data showed an interesting effect of the ECM component, which was different from our previous MEMA experiments where the ligand effects dominated the phenotype (Watson *et al*., 2018). In the 153LD1MY cells, fibronectin (FN1) and laminin strongly supported the binding and/or growth of cells, while collagens I, IV and 12A1 and vitronectin (VTN) showed more moderate binding (**Fig.1b**). Most of the other ECM molecules supported low levels of cellular binding and growth of the 153LD1MY cells. In contrast, for the A549 cells grown in low serum conditions, VTN showed the strongest binding and growth of cells, with collagens 12A1 and 1 showing moderate growth with COL4 and FN1 showed weak growth and binding. An expanded view of the data in bar graphs is shown in **Supplemental Figure 1a** and **1b**.

We concentrated our analysis on our the SASP-related ligands, as the ECM molecules used as substrate are aimed at providing an extracellular context that mimics general tissue microenvironments as opposed to SASP specific changes. For the 153LD1MY cells, we observed that 11 ligands significantly stimulated cell proliferation and thereby increased the number of cells per spot (FAS, CCL11, CCL16, CCL3, ANG, JAG1, FGF7, CSF3, TGFB2, TNFRSF10C, and EGF (**Fig.1b**); p<0.05 by Dunnett’s test). In contrast, we observed 31 ligands led to significantly lower numbers of cells per spot compared to control, including OSTP, IGFBP1, CSF1, IGFBP3, PIGF, CXCL12Beta, WNT10A, CXCL11, TNFRSF1A, CCL8, CXCL13, ICAM3, IL1B and CCL2, all of which showed a 40-fold or greater decrease in cell numbers compared to PBS control (p<0.05 by Dunnett’s test). For the A549 cells under low-serum conditions, the cell numbers were noticeably lower than for the 237LD1MY cells, presumably due to reduced growth rates. For A549, only TGF-β2 and KITLG resulted in significantly higher numbers of cells per spot than controls, although EGF also showed an almost 10x increase in cell number and borderline significance (p=0.18 by Dunnett’s test with corrections for multiple comparisons) (**Fig.1c**). In contrast, 15 ligands gave rise to significantly lower numbers of A549 cells compared to PBS control conditions, including MIF, IGFBP4, IL1B, CSF3, WNT10A, ICAM1, CXCL10, CCL16, and FGF2.

We next looked at EdU incorporation in the two cell lines to determine which conditions led to the highest degree of cell cycle activity. In the 153LD1MY cells, IL6, TGF-β1, TGF-β2, WNT7A, ANG and HGF led to a significantly higher proportion of EdU positive cells (p<0.05 by Dunnett’s test; **Fig.1d**), while 20 ligands significantly reduced the EdU positive proportion, including FGF7, IL13, CLC12 and FGF2. Since we have previously reported the effects of ligands from this assay on the EdU incorporation in the A549 cells grown in low-serum conditions (González-Gualda *et al*., 2022), we chose to interrogate the EdU incorporation as a function of ECM context (**Fig.1e**). In keeping with the results in **Fig.1c**, we saw the highest EdU incorporation in conditions that included COL1, FN1, COL4, COL2A1, COL3A1, and VTN cellular microenvironments.

### Validation of MEMA hits

Based on our preliminary findings, we selected several ligands to validate for their ability to impact the growth and survival of HMEC and lung cancer cells. For the HMEC cells, we focused on TGF-β1/2, HGF, ANG and IL-6, as these led to increased cell number and/or EdU incorporation in our MEMA experiments (**Fig.1b** and **1d**). Since we had multiple HMEC cell strains that differed in their degree of transformation, ranging from post-stasis to immortalized and transformed, we tested the ability of the selected ligands to impact cell growth and survival in several *in vitro* assays. We first tested the ability of the ligands to impact colony formation and growth by plating 500 cells into collagen-coated wells. As shown in **Fig.2a**, TGF-β1 and 2 lead to increases in colony formation and growth of the HMEC cells over 14 days compared to control cultures, but only in HMEC cultures derived from post-menopausal women, as TGF ligands had minimal impact on colony formation in the HMEC cells from young women. Quantification of crystal violet staining showed significant increases in cell numbers for TGF-β1 treated 237D1MYL cells compared to vehicle treated cells (**Fig.2b**). Unexpectedly, HGF had no impact on colony formation or cell growth in either HMEC strain, while IL-6 enhanced growth and colony formation in immortalized HMEC strains derived from both young and old donors. Finally, ANG showed modest colony formation stimulation, primarily in the HMEC from young donors.

**Figure 2.**
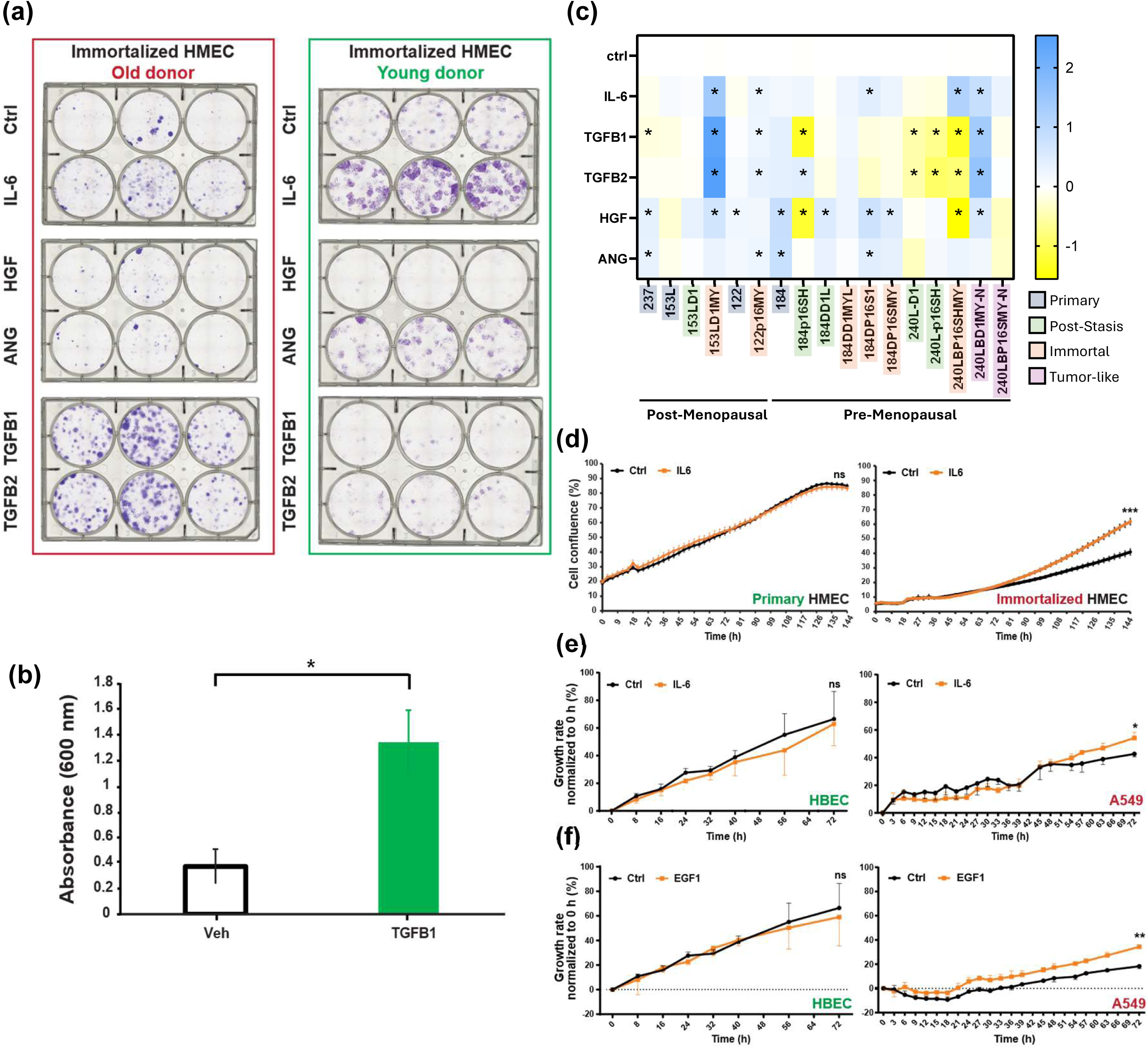
Targets identified from MEMA are validated using colony formation and live IncuCyte imaging assays. **(a)** HMEC were treated with different MEMA-identified hits with results for colony formation in cells immortalized cells derived from old vs. young donors showing significant enhancement of colony formation, particularly with TGF-β1 treatment in cells from aged donors. **(b)** Crystal violet quantification of colony numbers in control vs. TGF-β1 treated 237D1MYL HMEC cells. Triplicate samples were run and mean ± S.D. was plotted; * indicates p<0.05 by t-test. **(c)** Heat map summarizing the effects of IL-6, HGF, ANG TGF-β1, and TGF-β2 in 5 different HMEC cell strains tested. Strains are grouped by patient age as pre-menopausal (age 18-21) or post-menopausal (age 55-66), and individual cell lines names are color-coded by their mutational progression status; see **Sup. Table 1** for cell line details. * indicates significantly different (p<0.05) from control cells by ANOVA. **(d)** Example of IL-6 enhanced growth of primary cells (153L; left panel) vs. immortalized but not normal HMEC cells (153LD1MY; right panel) in live cell imaging experiments. All live cell imaging experiments were performed with four replicates and mean ± S.E.M. was plotted. **** indicates p<0.001 by t-test. **(e-f)** Validation of MEMA hits in normal human bronchial epithelial cells (HBECs) (left panels) or A549 (right panels). Data of 3-4 biological replicates are shown as mean ± S.E.M. * indicates p<0.05 and ** indicates p<0.01 by t-test.

We next performed live cell analysis of the growth for multiple HMEC cell strains in the presence of TGF-β, HGF, ANG and IL-6 over three days, as summarized in **Fig.2c**. For each ligand, the growth effect varied depending on both the progression status of the cell line as well as the age of the original patient. We observed that addition of IL-6 significantly increased the growth of immortalized HMEC cells from post-menopausal patients like 153LD1MY but not pre- or post-stasis cell from the same donor (**Fig.2c-d**). Results from pre-menopausal patients were similar, with immortalized and tumor-like lines showing increased growth,which reached significance in three out of five strains (**Fig. 2c**, right side). TGF-β1 and 2 only enhanced the growth of the immortalized HMECs that were derived from post-menopausal women, while inhibiting growth in most post-stasis and immortalized lines from younger women. One tumor-like line from a younger woman (240LBD1MY-N) did show significantly enhanced growth with TGF-β treatment, similar to the immortalized lines from older women, but the second immortalized line derived from this patient (240LBP16SMY-N) did not. HGF enhanced growth in most primary lines, as well as immortalized lines from older patients. In lines from younger patients, HGF effects were more variable, with one post-stasis line and one immortalized line from different patients strongly growth-inhibited by HGF while most others were enhanced nor not significantly affected. These data suggest that there may be fundamental differences between the cells of older and younger patients that affect how cells at different stages of tumorigenesis react to SASP signaling.

We performed similar validation studies in the A549 lung adenocarcinoma cells. Since we had previously done extensive validation of the effects of TGF-β in the A549 cells (Estela González-Gualda, 2022), we chose to study EGF, which stimulated cell number (and is related to AREG, which also impacted EdU positive proportion in the A549 cells). To understand the effect of IL-6 exposure in distinct cellular contexts, we also included it for testing in A549 cells, given the significant impact IL-6 had on HMEC cells, as shown above. In the live cell assays, both EGF and IL-6 treatment of A549 cells resulted in a small but significant increase in proliferation compared to control conditions (**Fig.2e-f**). In contrast, primary human bronchial epithelial cells (HBEC) showed no significant changes in growth when exposed to either of these ligands (**Fig.2e-f**). Of note, the increased proliferation rates of A549 cells upon EGF exposure were reversed to baseline levels by the concomitant treatment with an EGFR inhibitor (**Supplemental Figure 2**). As an internal control, we treated A549 cells with the same non-lethal dose of the EGFR inhibitor gefitinib and showed it had no significant impact on growth of control A549 cells at the concentrations tested (**Supplemental Figure 2**), which reinforces the functional role of EGF ligand in promoting A549 proliferation.

### Effects of Conditioned Medium on Cell Growth

We next sought to validate if conditioned medium isolated from senescent cell cultures *a)* could enhance cell growth; and *b)* contained the same growth-promoting factors we identified and validated from the MEMA experiments. To recapitulate the TME around A549 cells, we induced senescence in the human pulmonary fibroblast-adult (HPF-a) cells with chemotherapy (cisplatin; therapy-induced senescence, TIS) or radiation (radiation-induced senescence, RIS). We confirmed induction of senescence using various markers at both transcriptional and protein levels as well as senescence-associated β-galactosidase (SAβ-Gal) staining (**Supplemental Fig.3a-d**). We then collected medium (CM) from control, TIS- or RIS-treated HPF-a cultures and supplemented A549 cultures with the CM. We saw enhanced growth of A549 cells exposed to either TIS or RIS-conditioned medium from HPF-a (**Fig.3a**). We next used cytokine arrays to identify specific SASP factors that were overexpressed in the TIS and RIS CM compared to control CM. Remarkably, we found that EGF was significantly overexpressed in the senescent media from both TIS and RIS cultures compared to control CM (**Fig.3b-d**). Addition of gefitinib to the CM significantly impaired the growth of A549 cells treated with TIS or RIS CM, but not the control medium (**Fig.3e**) over a 4-day assay. In longer term colony formation assays, we observed that gefitinib inhibited the growth of both the control and TIS/RIS CM treated cultures as measured by both the number of colonies formed and crystal violet staining and spectrophotometer measurements of optical density (**Fig.3f-h** and **Supplemental Fig. 3h-j**).

**Figure 3.**
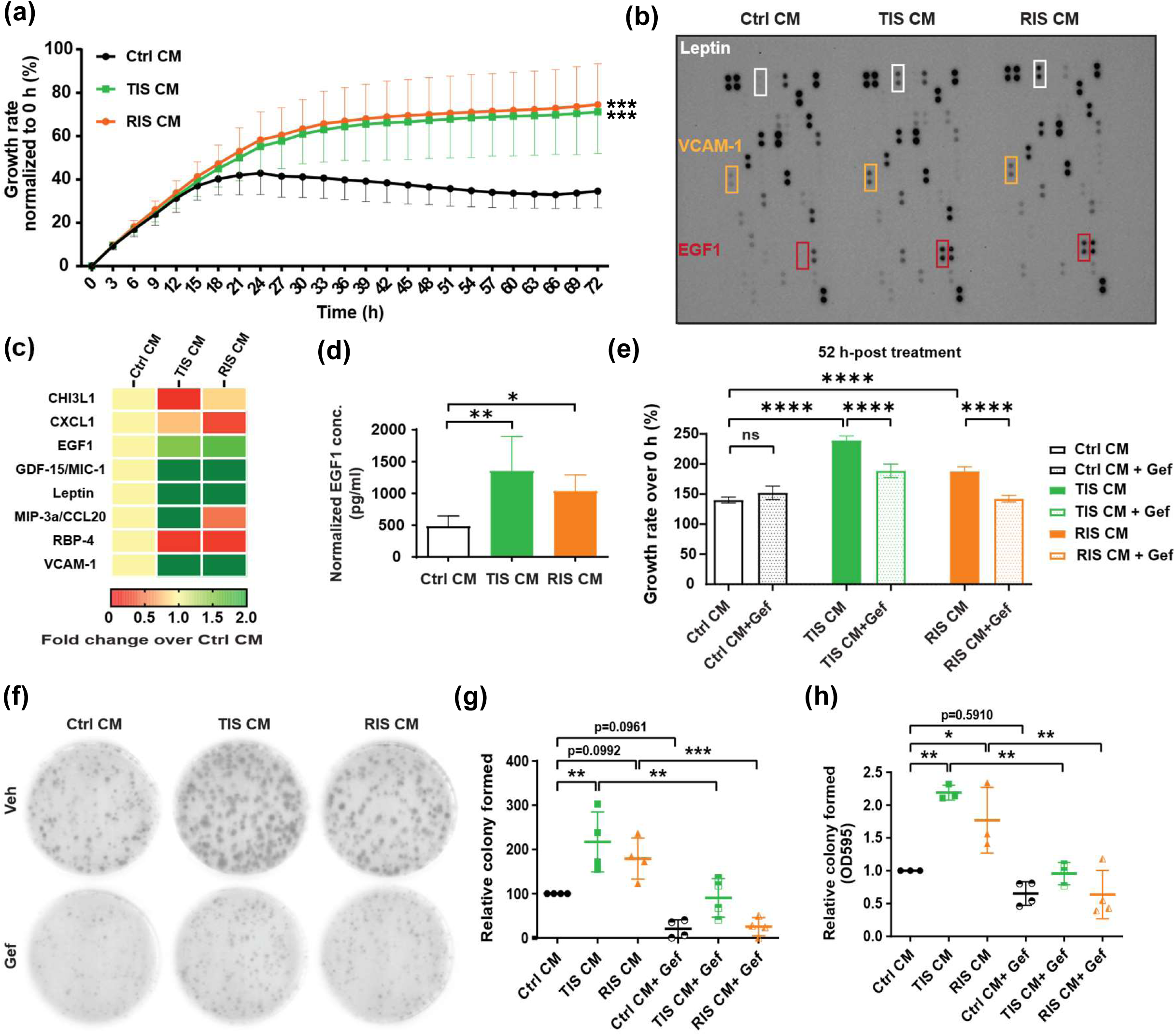
Senescent conditioned media (CM) promote growth of A549 cells through SASP factor EGF1. **(a)** Growth promotion of A549 cells by senescent CM compared to control CM of human pulmonary fibroblasts – adult (HPF-a). Cells were seeded at a density of 25,000 cells per well in a 24-well plate the day before treated with indicated CM. Real-time cell growth was monitored using IncuCyte® SX5 Live-Cell Analysis Instrument (Sartorius AG) for 3-5 days. Cell growth was analysed using the IncuCyte® software and manifested as percentage over the baseline confluence. Data of 6 biological replicates are shown as mean ± SD. **indicates significant difference (p<0.01) from cells treated with Ctrl CM as determined by ANOVA analysis. **(b-c)** Secretome analysis of CM from control or senescent HPF-a. CM were harvested after 72-h incubation with corresponding HPF-a and the components were evaluated using Proteome Profiler Human XL Cytokine Array kit (#ARY022B, R&D Systems). VCAM-1, vascular cell adhesion molecule 1. The original blots **(b)** and quantification results are shown as heatmap **(c)**. **(d)** Validation of EGF1 level in CM with ELISA. The CM were harvested and subjected to ELISA (#DEG00, R&D Systems) for determining the amounts of EGF1 with normalization based on cell contents. Data of 6 biological replicates are shown as mean ± SD. *p < 0.05; ** p < 0.01 as determined by ANOVA analysis. ns, not statistically significant. **(e)** Reversion of CM-enhanced cell growth by EGF receptor blocker gefitinib (Gef). A549 cells were cotreated with CM from control or senescent HPF-a with or without gefitinib (2.5 μM) before subjected to IncuCyte® SX5 Live-Cell Analysis Instrument (Sartorius AG) for 3-5 days. Representative data of 3 independent biological repeats are shown as mean ± SD. **** p < 0.0001 as determined by ANOVA analysis. **(f-h)** Reversion of CM-enhanced colony formation by EGF receptor blocker gefitinib (Gef). A549 cells were seeded at a density of 750 cells per well in a 6-well plate the day before treatment with designated HPF-a CM and/or gefitinib for 15 days. Treatment was refreshed every 2-3 days. Colonies were stained with crystal violet and images were scanned for quantification as described in the methodology. Representative images **(f)**, colony count **(g)**, and OD595 **(h)** of 3-4 independent biological replicates are shown as mean ± SD. *p < 0.05; ** p < 0.01; *** p < 0.001 as determined by ANOVA analysis.

We performed similar experiments in the 237LD1MY HMEC cells. We used sub-lethal doses of cisplatin, doxorubicin, or paclitaxel to induce senescence, which was confirmed by increased SAβ-Gal staining and a decrease in EdU incorporation (**Supplemental Fig.4a, b**). As observed in the A549 cells, CM collected from the senescent cell cultures significantly enhanced cell growth compared to control CM (**Fig.4a**). We again utilized the cytokine arrays to assess differential growth factor expression between the TIS and control CM cultures (**Fig.4b**). The arrays lacked TGF-β for testing, but we did observe up-regulation of IL6, particularly in the paclitaxel CM (**Fig.4b**), as well as other factors like CXCL5, OPN, and IL8. We also tested the CM for TGF-β1 by ELISA and found that it was elevated in the CM from cisplatin- and doxorubicin-treated cells relative to control CM, but not in the paclitaxel-treated CM (**Fig.4c**). We next tested the ability of the TGFBRI inhibitor galunisertib to block the growth-enhancing effects of cisplatin CM. We used a dose of 2 µM galunisertib, which was the highest dose that did not impact 237D1MYL growth in a 3-day growth assay (**Supplemental Fig.5**). Galunisertib treatment of the cells growing in control CM had no impact on cell numbers, but resulted in a small decrease in cell number in the cells growing in cisplatin CM (**Fig.4d**), although this failed to reach significance. This suggests that other factors in addition to TGF-β might also be involved in the observed phenotype and/or that galunisertib efficiency to block the specific TGFBR is suboptimal.

**Figure 4.**
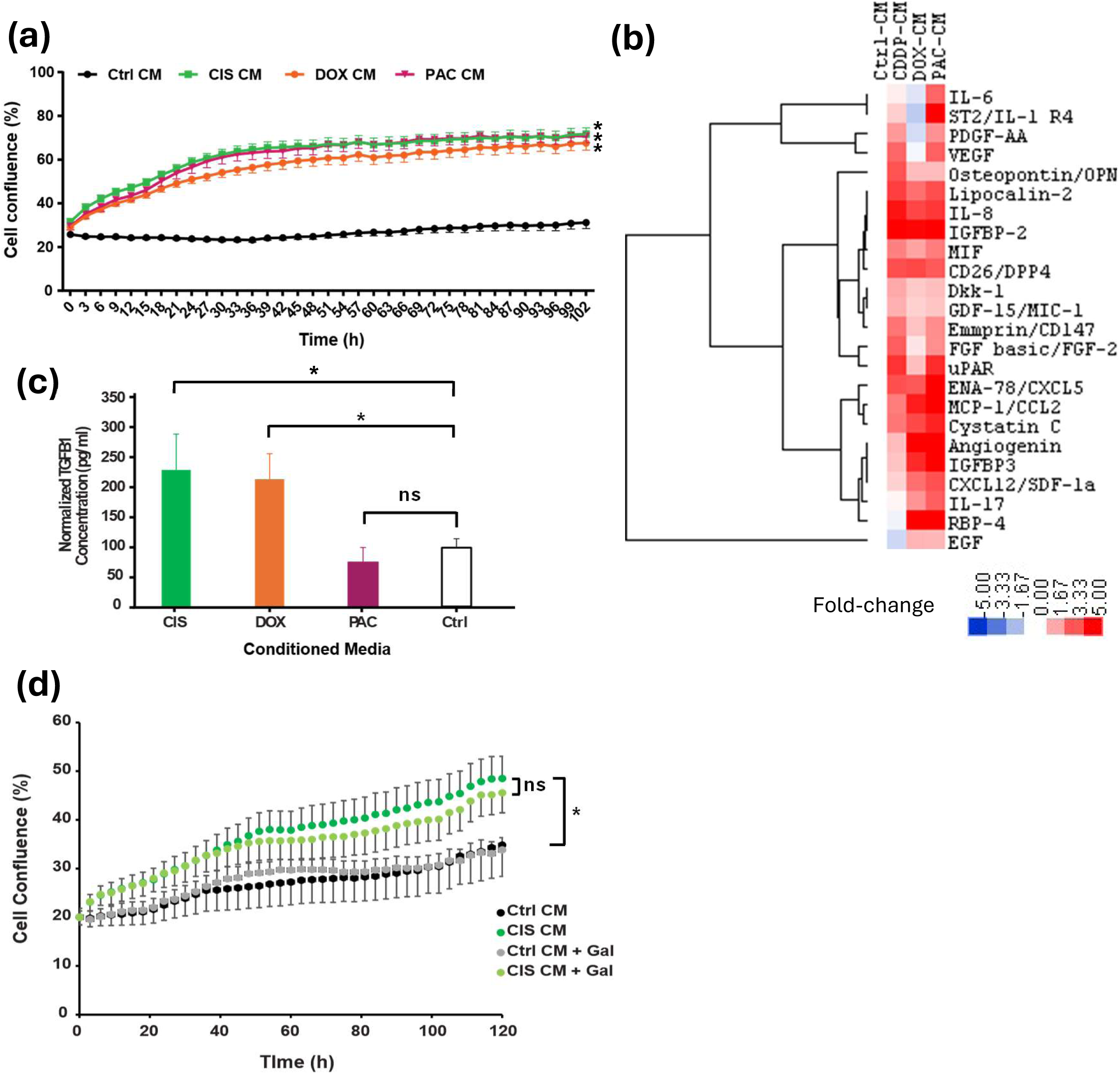
Senescent conditioned media (CM) promote growth of HMEC cells. (**a**) Growth promotion of HMEC by senescent CM induced by different chemotherapeutic agents compared to control CM of HMEC. Representative data of 3 independent biological repeats are shown as mean ± SD. CIS, cisplatin; DOX, doxorubicin; PAC, paclitaxel. * indicates significant difference (p<0.05) in growth curves as determined by ANOVA analysis. (**b**) Heatmap summarizing the secretome analysis of CM from control or senescent HMEC. CM were harvested after 24-h incubation with corresponding senescent HMEC cultures and the components were evaluated using Proteome Profiler Human XL Cytokine Array kit (#ARY022B, R&D Systems). (**c**) ELISA analysis of TGF-β1 levels from conditioned medium from cisplatin, doxorubicin, paclitaxel, or PBS control treated conditions. * indicates significant difference in levels compared to control as determined by t-test. ns, not statistically significant. (**d**) Enhanced growth of 237D1MYL HMEC cells by cisplatin conditioned medium is partly inhibited by addition of 2 µM galunisertib, which has no effect on the growth of cells under control conditions. * indicates significant enhancement of growth in cells grown in cisplatin-conditioned medium compared to controls.

**Figure 5.**
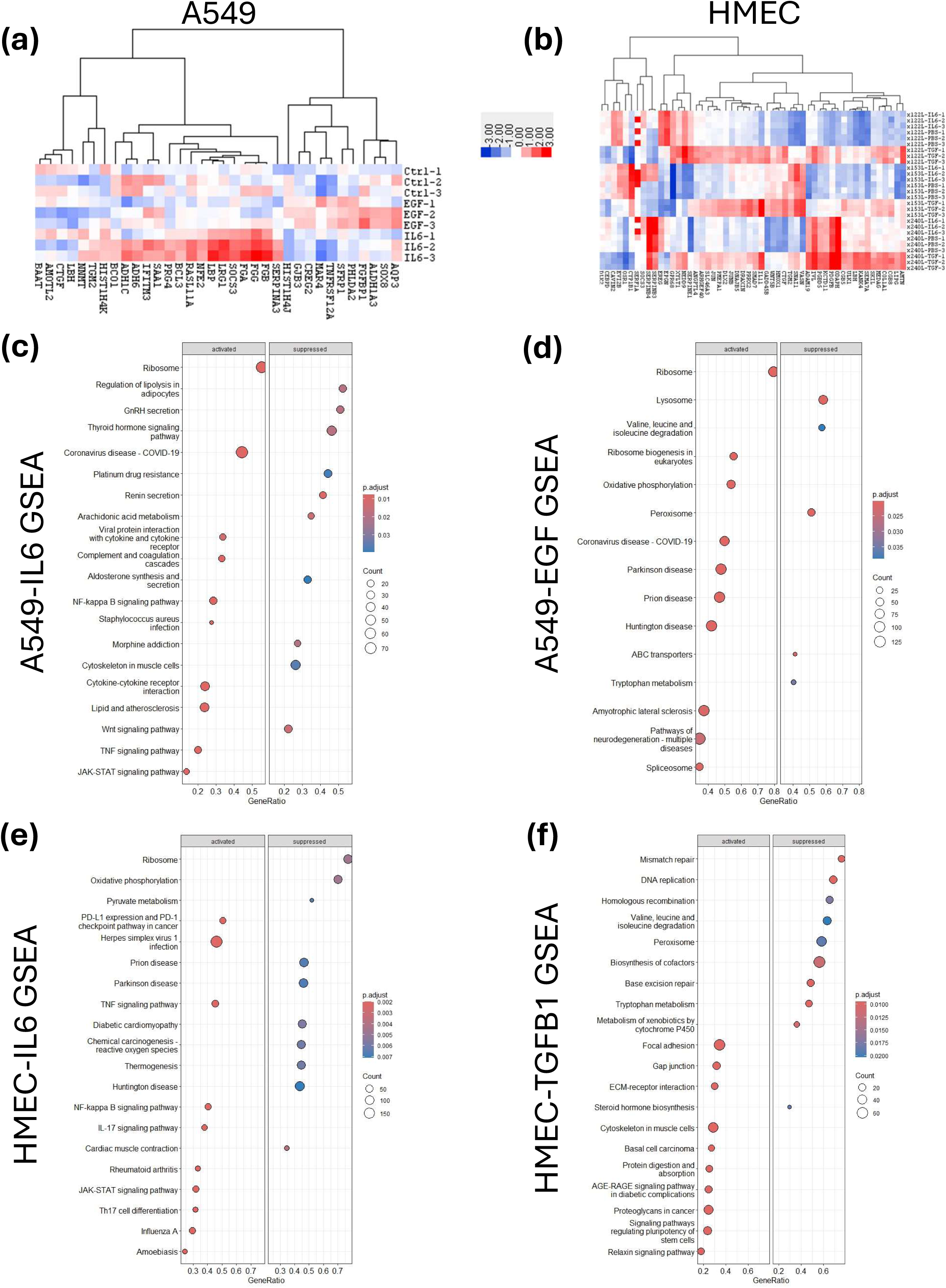
Context-dependent gene expression profiles in response to SASP factors. Cells were treated with corresponding SASP factors for 6 h before subjected to RNA extraction and subsequent RNAseq. Heatmaps for lung (**a**) and HMEC (**b**) cells showing the genes with the largest differential expression as a result of treatment with the indicated ligands. GSEA analysis using KEGG databases show significantly altered pathways and processes in A549 cells treated with IL-6 (**c**) and EGF (**d**). GSEA analysis using KEGG databases show significantly altered pathways and processes in 122L, 153L, or 240L HMEC cell lines treated with IL-6 (**e**) or TGF-β1 (**f**). Dot size indicates number of genes within a given set and colour indicated adjusted p-value.

### RNAseq Analysis of SASP Factor Impact

We next sought to understand how exposure to the various SASP factors was altering gene expression. We performed RNAseq analysis on A549 lung cancer cells and 122L and 153L (HMEC cultures derived from post-menopausal women), and 240L (HMEC cells derived from an 18-year-old woman) that had been treated with EGF (A549 only), TGF-β1 (HMEC only), or IL-6 (both A549 and HMEC) for six hours. In the A549 cells, we identified 26 transcripts that were at least 2-fold more highly expressed in the ligand treated cells compared to controls, and 10 transcripts that were down-regulated by 2-fold or more (**Fig.5a**). The biggest changes in expression were TNFRSF12A (EGF-treated) and LRG1 (IL-6 treated), showing 3.4- and 5-fold increases in expression following treatment respectively. In the HMEC cells, we again looked for genes that showed at least 2-fold changes in expression on average across the three HMEC cell strains treated with ligand compared to untreated cells. In total, there were only 52 genes that showed at least a 2-fold change in expression, and again the majority of them (48) were in the TGF-β treated cells. Only 8 genes showed down-regulation in response to SASP ligand, with the rest of the genes were upregulated in response to treatment. We show representative heatmaps of all the differentially expressed genes in A549 (**Fig.5a**) and the top 50 genes in the HMEC lines (**Fig.5b**).

Next, we performed differential gene expression analysis followed by gene set enrichment analysis (GSEA) using Gene Ontology (GO) and KEGG pathway data to identify pathways and processes enriched as a result of SASP ligand treatment. The GO GSEA analysis for A549 showed that IL-6 treatment induced cytokine response pathways and EGF treatment was most strongly associated with activation of ribosomes and rRNA processing (**Supplemental Fig.6a-b).** The GO GSEA analysis for the HMEC cells treated with TGF-β showed induction of wound healing programs, along with increased cellular motility and tissue morphogenesis, and treatment of the HMEC cells with IL-6 elicited an immune/cytokine associated response (**Supplemental Fig.6c-d**). For the KEGG analyses, treatment of A549 with IL-6 activated JAK-STAT and NF-kappa B signaling pathways and down-regulated platinum drug resistance (**Fig.5c**). Interestingly, EGF treatment of A549 activated multiple pathways associated with ribosomes and ribosome biogenesis as well as oxidative phosphorylation (**Fig.5d**). For the KEGG analysis of HMEC cells, we found that IL-6 again activated JAK-STAT and NF-kappa B (a well know SASP driver) signaling, as well as upregulation of PDL1 and PD-1 immune checkpoint expression (**Fig.5e**). The TGF-β treatment of HMEC cells induced pathways associated with pluripotency and stem cells and basal cell carcinoma, while downregulating several DNA repair pathways (**Fig.5f**).

### Targeting SASP drivers in vivo

We next sought to understand if these SASP factors were active *in vivo*. Since the specific HMEC lines we tested have not been reported to grow as xenografts, we focused our attention on the A549 cell line. As mentioned above, we have previously examined the impact of inhibiting TGF-β in studies of A549 cells, including *in vivo* (González-Gualda *et al*., 2022), so we again focused on the effects of EGF inhibition in these cells. We implanted A549 cells expressing a luciferase reporter into immune-compromised mice along with control and senescent fibroblasts (see Materials & Methods). When the tumors reached 100 mm^3^, we began treating the mice with 50 mg/kg of the EGF inhibitor gefitinib (**Fig.6a**). Treatment continued for 3 weeks and tumour size was measured by calipers and by bioluminescence. At the start of treatment, we did not see any growth enhancement with the inclusion of senescent fibroblasts compared to normal fibroblasts by either tumour volume or luminescence readings, although inclusion of normal fibroblasts significantly increased growth of the A549 tumors compared to controls with only A549 implanted, probably due to the contribution of some fibroblast proliferation to the tumor burden (**Fig.6b, c**). By 19 days however, the vehicle-treated tumours that contained senescent fibroblasts were significantly larger than the tumors that contained control fibroblasts (**Fig.6e**, p=0.0429). Gefitinib treatment showed a trend towards inhibiting the growth of the tumours that included senescent fibroblasts (**Fig.6d, e**), although it just failed to reach significance (p=0.0744). In contrast, the tumours harbouring normal control fibroblasts showed no signs of inhibition with gefitinib.

**Figure 6.**
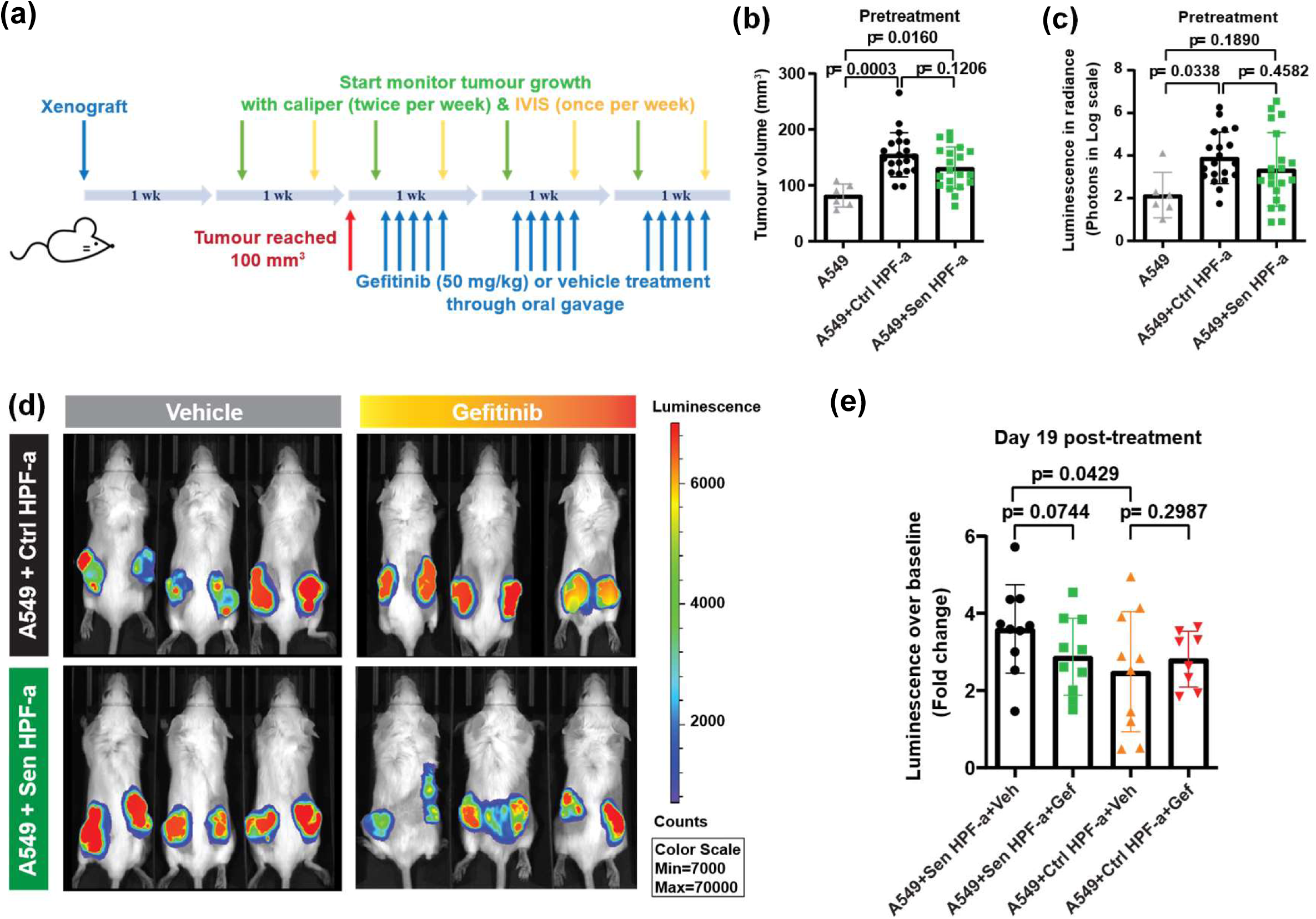
Systematic *in vivo* blockade of EGF receptor alleviates the tumour growth promoted by senescent microenvironment. (**a**) Work scheme of mice model bearing xenografts with A549-Luc2 cells alone or co-injected with control or senescent HPF-a. IVIS, *in vivo* imaging system. (**b-c**) Tumor measures prior to treatment with gefitinib. Total tumor volumes were calculated based on calliper measurement (**b**) whilst the tumor growth attributed to A549-Luc2 cells was evaluated using D-Luciferin and IVIS Spectrum Xenogen imaging system (Caliper Life Sciences) and analysed with Living Image^®^ software (ver 4.5.5) (**c**). p values are determined using ANOVA analysis. (**d-e**) Representative images (**d**) and quantification results (**e**) of tumor development after completing gefitinib treating regimen. Data are shown as mean ± SD with each data point representing per tumor. One-sided t-test was used to evaluate statistically the effect of senescent HPF-a on promoting the tumour growth and gefitinib on reversing such enhancement.

## Discussion

SASP has long been implicated in driving tumor growth, but it has been difficult to systematically address due to its complexity and heterogeneity. We reasoned that SASP would be an excellent test of our MEMA platform to identify specific SASP factors that could impact cancer cell growth and survival. MEMA allow for systematic analysis of multiple factors and combinations of them with/without other proteins (e.g. ECM proteins) within a single assay. We tailored the MEMA for this study to include multiple SASP factors in order to assess a broad collection of proteins that are associated with the senescence-associated phenotype for their impact on cell growth and survival of both breast and lung pre-cancer and cancer cells, respectively. We found that multiple SASP factors have the potential to drive growth of both lung cancer and pre-neoplastic breast cancer cells. Interestingly, there was significant overlap among the factors that enhanced cell growth and survival of the lung and breast cells. For example, the MEMA studies identified both TGF-β and IL-6 as enhancers of the growth and survival of lung cancer and HMEC cells, and both have long been associated with SASP.

IL-6 has been implicated in creating a pro-tumorigenic microenvironment (reviewed in (Wu *et al*, 2025)) that can drive early tumorigenesis. IL-6 signalling is usually confined to immune cells and the effects on tumour cells is indirect (Wu *et al*., 2025), since epithelial cells typically lack expression of the IL-6 receptor. Thus, IL-6 is thought to act on immune cells which then secrete additional factors that can aid in the growth of early pre-neoplastic lesions. However, in both our HMEC and lung cancer studies, we found that IL-6 could directly act on the pre-neoplastic epithelial cells. It is unlikely that the concentration of IL-6 is so high as to activate off-target receptors, as the concentration we used was only 0.6 ng/ml, which is the concentration utilized for on-target studies of IL-6 in immune cells. Instead, recent reports suggest that IL-6 can work in cells that lack IL-6 receptor, as soluble IL-6R can bind to IL-6 and together activate the gp130 co-receptor, resulting in downstream signalling and creation of a pro-inflammatory microenvironment (Tanaka *et al*, 2014). Interestingly, there are reports that bronchial epithelial cells can express IL-6 (Takizawa *et al*, 1996), and that IL-6 can act as a paracrine growth in breast cancer cell lines, which can upregulate both the soluble IL-6R and membrane bound IL-6R (Chiu *et al*, 1996). Thus, it is possible that IL-6R is upregulated in our cells. In fact, it was reported that IL-6 promotes growth and EMT in CD133+ NSCLC cell lines, including A549 cells (Lee *et al*, 2016). IL-6R signaling can promote stemness, growth and inflammatory responses in A549 cells and hence interventions like the IL-6R-blocking antibody tocilizumab have been used to investigate anti-cancer and anti-inflammatory effects on these cells (Kim *et al*, 2015). Interestingly, IL-6R expression appears to be a function of the degree of transformation, as the normal lung and mammary epithelial cells we tested were non-responsive to IL-6, with growth enhancement only occurring in the cancerous A549 cells and the post-stasis, immortalized HMEC cultures. Additional studies are underway to characterize expression of IL-6R and gp130 and the pathways activated in response to IL-6 in both the lung and HMEC cells.

One of the most striking differences we observed was the differential ability of TGF-β to enhance growth and survival of immortalized HMEC cultures derived from post-menopausal but not pre-menopausal women. Previous studies have shown that aging shifts the balance of luminal/myoepithelial lineages and changes multipotent progenitors, resulting in a basal differentiation state bias and an increased potential for malignant transformation {Garbe, 2012 #299}. TGF-β has typically been reported to have antagonistic effects on tumorigenesis, but some investigators have suggested that TGF-β may promote progression and an immunosuppressive microenvironment during tumorigenesis (reviewed in (Massague & Sheppard, 2023)). For example, some studies demonstrated that TGF-β may act as a tumour suppressor in the early tumour microenvironment (Pickup *et al*, 2013). In contrast, other investigators have shown pro-tumorigenic effects of TGF-β, including in pre-neoplastic breast cells, where TGF induction can drive epithelial to mesenchymal transition (EMT), resulting in a more aggressive phenotype that further drives progression (Andarawewa *et al*, 2007). We speculate that the changes to luminal/myoepithelial lineages and/or multipotent progenitors in post-menopausal mammary glands may result in acquisition of a TGF-β responsive state. We are currently investigating the effects of TGF-β on differentiation states and progenitor cells to better understand its importance in early mammary tumorigenesis in young versus older women.

Similarly, TGF has been implicated in driving EMT in lung cancer (Ramundo *et al*, 2023), but this is thought to occur later in progression as opposed to during early tumorigenesis. These studies align with the data we previously generated, as we found that therapy-induced senescence of cancer and stromal cells in response to platinum therapy results in a pro-tumorigenic SASP enriched in TGF-β ligands that activate TGFBR1 signalling and activation of AKT/mTOR pathway in recipient lung cancer cells, boosting tumor progression and relapse, and significantly reducing survival (González-Gualda *et al*., 2022).

TGF-β induced wound healing and/or acquisition of a stem cell phenotype, as per GSEA analyses, in both the HMEC primary cells we used. Interestingly, senescence and SASP have been implicated in creating a “stemness” phenotype that reprograms senescent cancer cells (TIS) into self-renewing, tumour-initiating stem-like cells that are associated with highly aggressive disease (Milanovic *et al*, 2018). Our data suggests that TGF-β may be one of the SASP factors that is capable of driving this stemness signature. Similarly, senescence has been implicated in normal wound healing, as it mediates changes in matrix production and vascularization (Wang *et al*, 2024). With early expression profiling studies, it was recognized that many cancers had expression signatures that showed a high degree of similarity to wound signatures (Chang *et al*, 2004), but the relationship between senescence and a wound healing signature in cancer remains incompletely characterized (Pulido *et al*, 2021). Interestingly, IL-6 has also been implicated in wound healing and SASP (Pulido *et al*., 2021), demonstrating that multiple factors associated with SASP are likely capable of driving acquisition of both the wound healing and stem cell signatures in cancer.

EGF elicited a growth enhancement in the A549 cells *in vitro*, which we further validated *in vivo*. The effects on growth of A549 xenografts was significantly enhanced by the inclusion of senescent fibroblasts, which was partially inhibited by the addition of the EGFR inhibitor gefitinib. Although this result failed to reach statistical significance there was a clear trend towards a proliferative effect. As mentioned earlier, we have also tested the effects of TGF-β inhibitors on the growth of A549 tumors in mice (González-Gualda *et al*., 2022), where tumor volume was significantly restored but not completely to baseline levels. This likely reflects the multifactorial and dynamic nature of the SASP, and thus a combination of several factors at different longitudinal timelines post-therapy may be involved in promoting tumor growth and other protumorigenic activities. Based on these results, it is tempting to propose that the use of senolytics to target senescent cells to remove all SASP and restore the microenvironment to a non-inflammatory state might be an effective treatment strategy. However, given the toxicity of many senolytics like navitoclax and dasatinib+quercetin, we believe a better approach is likely to target specific SASP drivers that are identified in studies like ours to prevent tumorigenesis.

The impact of the SASP factors discussed above was limited to pre-neoplastic or fully transformed cells, having no impact on cell growth or survival in more normal breast and lung cell populations. This supports the notion that efficient tumorigenesis depends on at least two factors: the presence of mutated, pre-neoplastic cells, and an altered, pro-tumorigenic microenvironment. Either one of these factors on its own is unlikely to result in tumorigenesis, as a healthy microenvironment appears capable of restricting pre-neoplastic cell growth and progression, while normal cells appear to be non-responsive to the tumour promoting effects of an inflammatory microenvironment. In contrast, we did witness growth-enhancing effects on primary HMEC cells by HGF and ANG, which had either mixed (HGF) or little (ANG) effects on more-progressed cells. This further indicates that the specific makeup of the SASP in a given tissue microenvironment is an essential consideration in tumorigenesis and progression.

In summary, we have utilized our MEMA platform to perform a systematic, unbiased screen of putative SASP factors for their ability to drive growth and survival in pre-neoplastic and fully cancerous breast and lung cells. We find that some of the same factors have the potential to drive these processes in both tumour types. Furthermore, we find that pro-tumorigenic SASP factors, such as EGF1, are present in conditioned media collected from senescent cells, and that this conditioned media significantly enhances the growth and survival of pre-neoplastic and cancer cells, but not normal cells. Our expression profiling of the response of cells shows early induction of multiple cellular programs associated with increased proliferation (e.g., increased protein synthesis) as well as acquisition of stem-like and wound healing signatures. Finally, we show that these SASP factors are likely to functional *in vivo*, as targeting factors like TGF-β and EGF in lung cancers containing senescent cells appears to inhibit tumour cell growth. However, the impact of the SASP is likely multifactorial, as targeting individual SASP factors does not restore growth to baseline, suggesting that targeted inhibition of combinations of key SASP factors like those identified by our MEMA studies might be the optimal therapeutic approach. In short, our MEMA studies have identified key SASP factors that drive tumorigenesis and provides strategies for either targeted removal of senescent cells or inhibition of key SASP factors as a potential therapeutic approach to treating early breast and lung cancer lesions.

## Acknowledgments

JEK and RH’s work on this project was supported in part by the Cancer Early Detection Advanced Research Center (CEDAR) OHSU Knight Cancer Institute, Project Award funding (grant 2018-CRUK-OHSU-003). Further work in the Korkola lab was supported by an NCI R21 award (grant 1R21CA286419-01A1). The Muñoz-Espín’s laboratory was supported by the Cancer Research UK (CRUK) Cambridge Centre Early Detection Programme (RG86786), by a CRUK Programme Foundation Award (C62187/A29760), by a CRUK Early Detection OHSU Project Award (C62187/A26989), by a Medical Research Council (MRC) New Investigator Research Grant (NIRG) (MR/R000530/1) and by a Darley/Sands Downing College Fellowship (G109261). H-LO was funded by a CRUK Early Detection OHSU Project Award (C62187/A26989).

## Notes

Conflict of Interest statement: Dr. Korkola is a co-founder and stock holder in Convergent Genomics. The work described in this manuscript is not related to any work being performed at Convergent Genomics. The other authors have stated that they have no conflicts of interest in connection with this article.

### Competing Interest Statement

The authors have declared no competing interest.

