## Supplemental for "Identification of senescence-associated drivers of tumour growth and progression using a novel microarray platform"

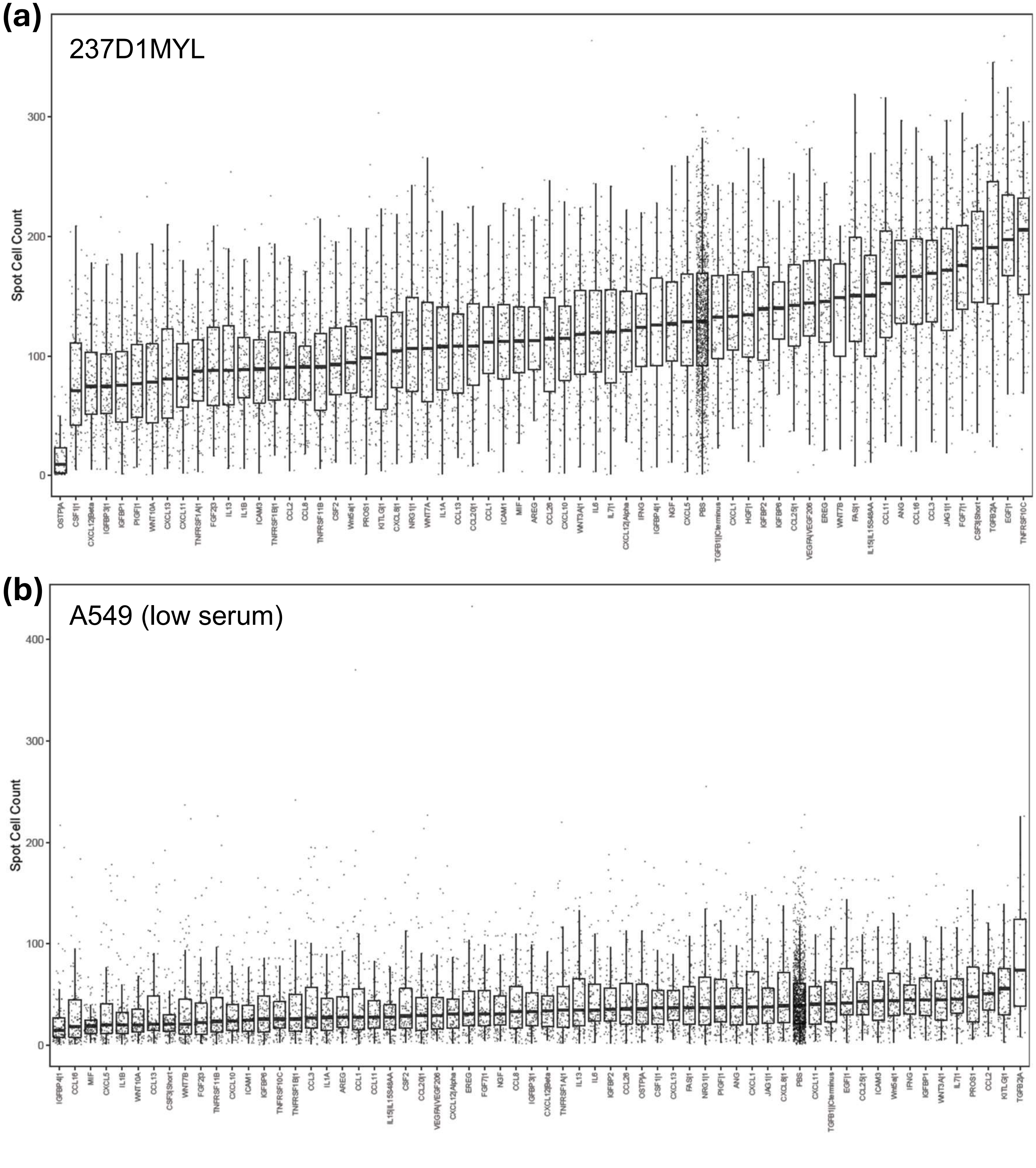


**Supplemental Figure 1.** Spot cell count for 237D1MYL (**A**) and A549 (low serum; **B**) resulting from exposure to different ligands.

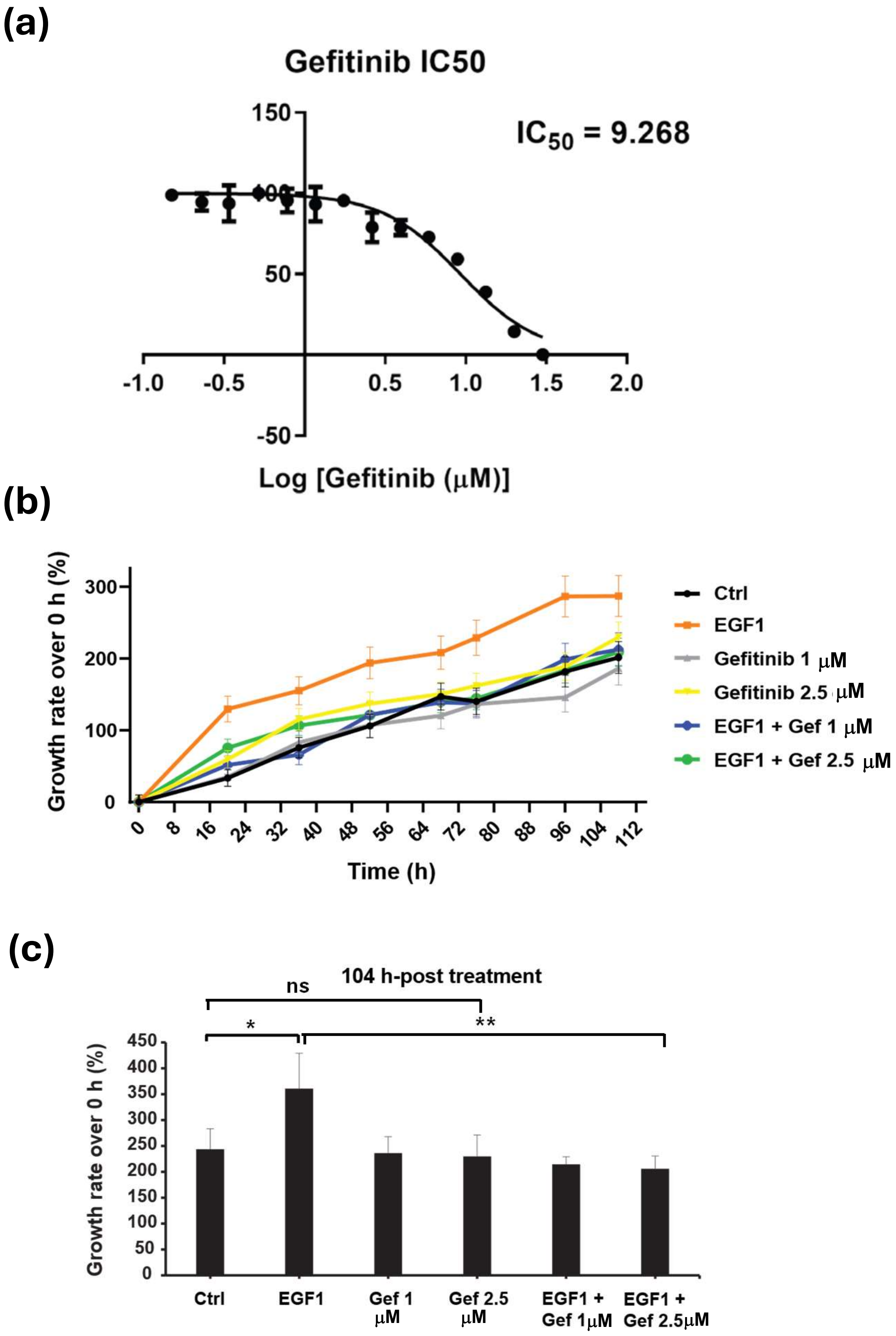


**Supplemental Figure 2.** Effect of the EGFR inhibitor gefitinib on A549 cell growth. (**A**) A549 cells show limited response to gefitinib over the course of 3 days, with an $IC_{50}$ of 9.268 µM. (**B**) A549 cells treated with EGF show enhanced growth that is reversed by treatment with gefitinib (1 or 2.5 µM); in contrast, these doses of gefitinib have no significant effect on A549 in the absence of EGF. Representative data of 3 independent biological repeats are shown as mean ± SD. (**C**) The growth rate at 104 h-post treatment are shown in bar chart. One-way ANOVA + Tukey HSD test.

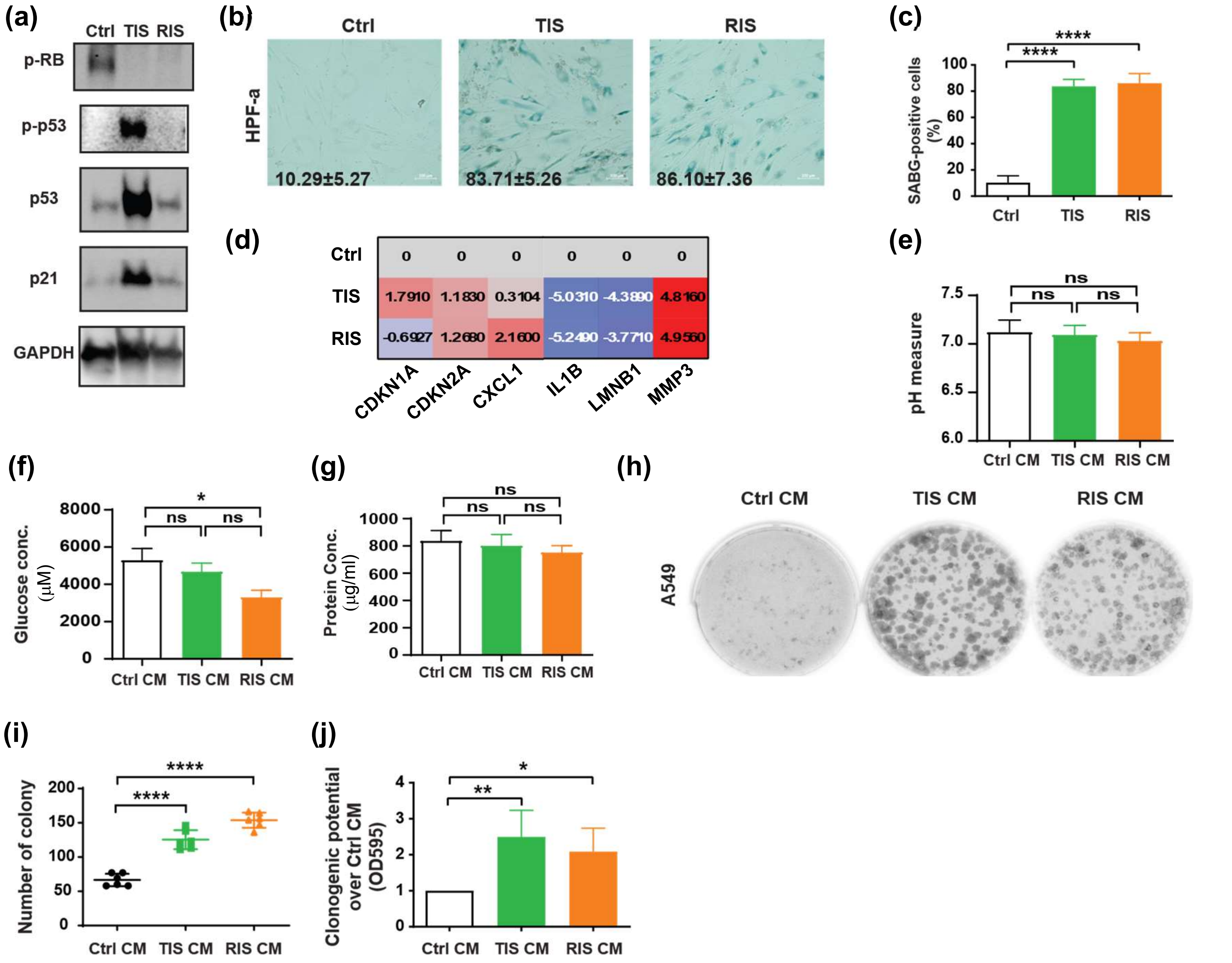


**Supplemental Figure 3.** Characterization of cellular senescence in HPF-a upon treatment with cisplatin or ionising radiation (IR) and corresponding conditioned media (CM). (a) Protein expression of senescence markers in normal HPF-a or senescent HPF-a induced with cisplatin (TIS, therapy-induced senescence) or IR (RIS, radiation-induced senescence). (b-c) Representative images (b) and quantification (c) of HPF-a stained for SA-b-Gal. A549 cells were untreated (Ctrl) or treated with cisplatin or IR for senescence induction. Cells were fixed on Day 10 after senescence induction and subjected to SA-b-Gal staining using Senescence b-Galactosidase Staining Kit (#9860, Cell Signaling). Images were taken using Zeiss Axio Observer 7 microscope and ZEN Blue software (ver 3.1). Scale bar, 100 mm. Data are shown as mean ± SD from 5 biological replicates. (d) Transcriptional expression of senescence-associated genes detected using quantitative real-time PCR. RNA of A549 cells was extracted on Day 10 after senescence induction and subjected to RT-qPCR. The heatmap are shown as mean fold change in $\log_2$ compared to Ctrl from 3-5 biological replicates. (e-g) Characterization of 72-h CM from Ctrl, TIS, or RIS HPF-a in terms of the pH value (e), glucose concentration (f), and protein levels (g). (h-j) Growth promotion of A549 cells by senescent CM. Colony formation assay was performed to evaluate the impact of senescent CM on the growth of A549. Cells were seeded at a density of 750 cells per well of 6-well plate one day before the treatment. CM treatment was refreshed every 2-3 days throughout the 15-day assay. Representative images (h) and quantification (i-j) of colony formed from 4 biological replicates are shown as mean ± SD. *$p < 0.05$; ** $p < 0.01$; **** $p < 0.0001$; ns, not significant.

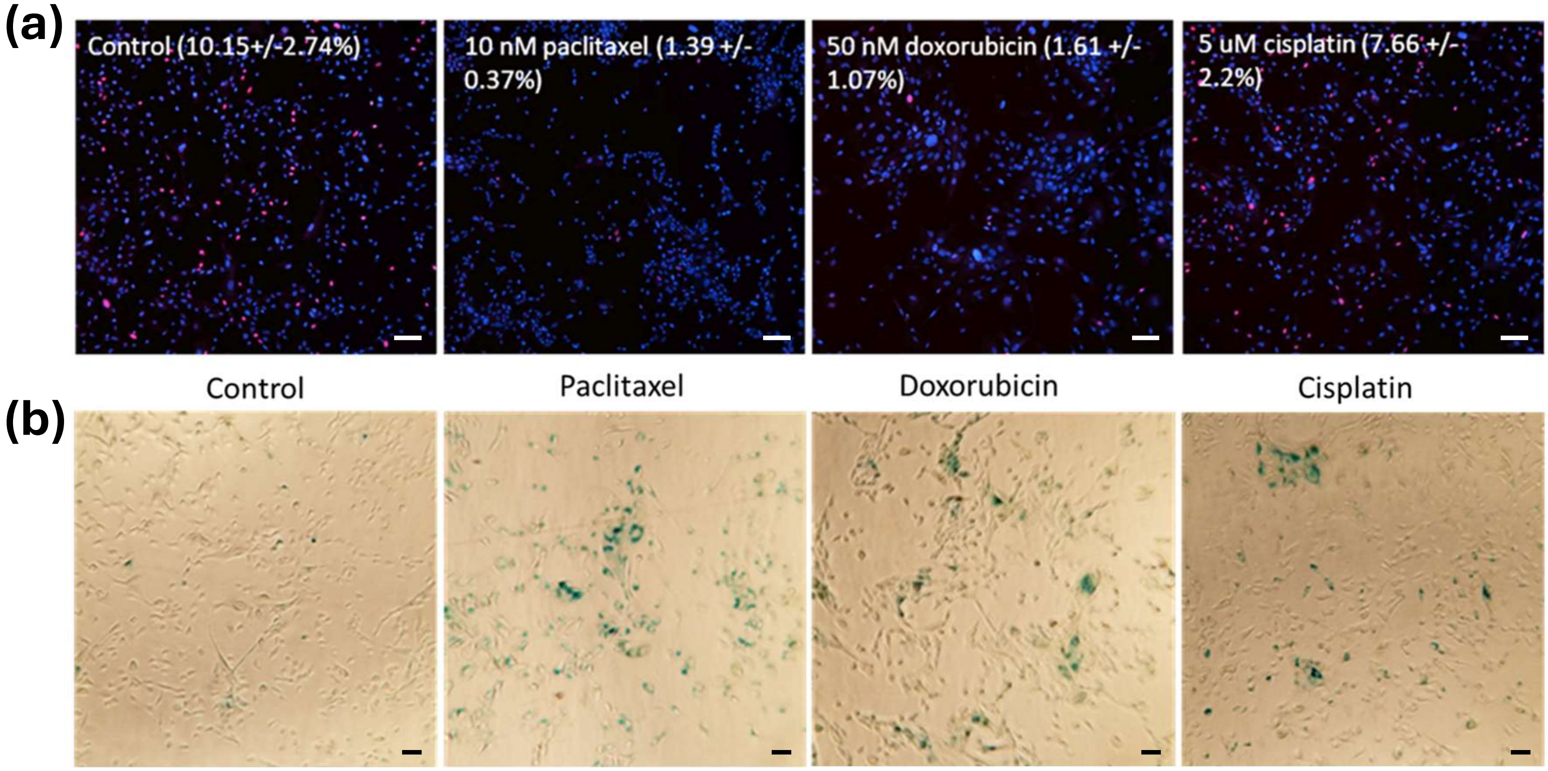


**Supplemental Figure 4.** Induction of senescence through treatment with chemotherapeutic agents. **A**. Induction of senescence with chemotherapy treatment in the HMEC cell strain 237D1MYL as measured by loss of EdU incorporation or **B.** SABgal staining (**B**). Values in **A** represent percentage of cells positive for EdU incorporation, +/- S.D. from triplicate wells. Scale bar in A is 100 μm and scale bar in B is 50 μm.

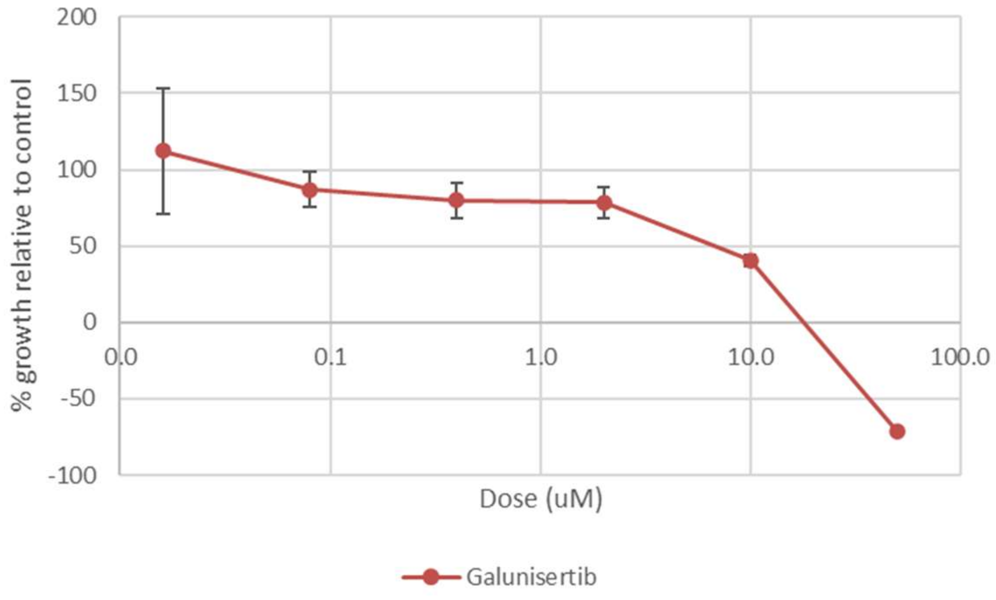


**Supplemental Figure 5.** Effect of the TGFBR inhibitor galunisertib on HMEC 237D1MYL cell growth.

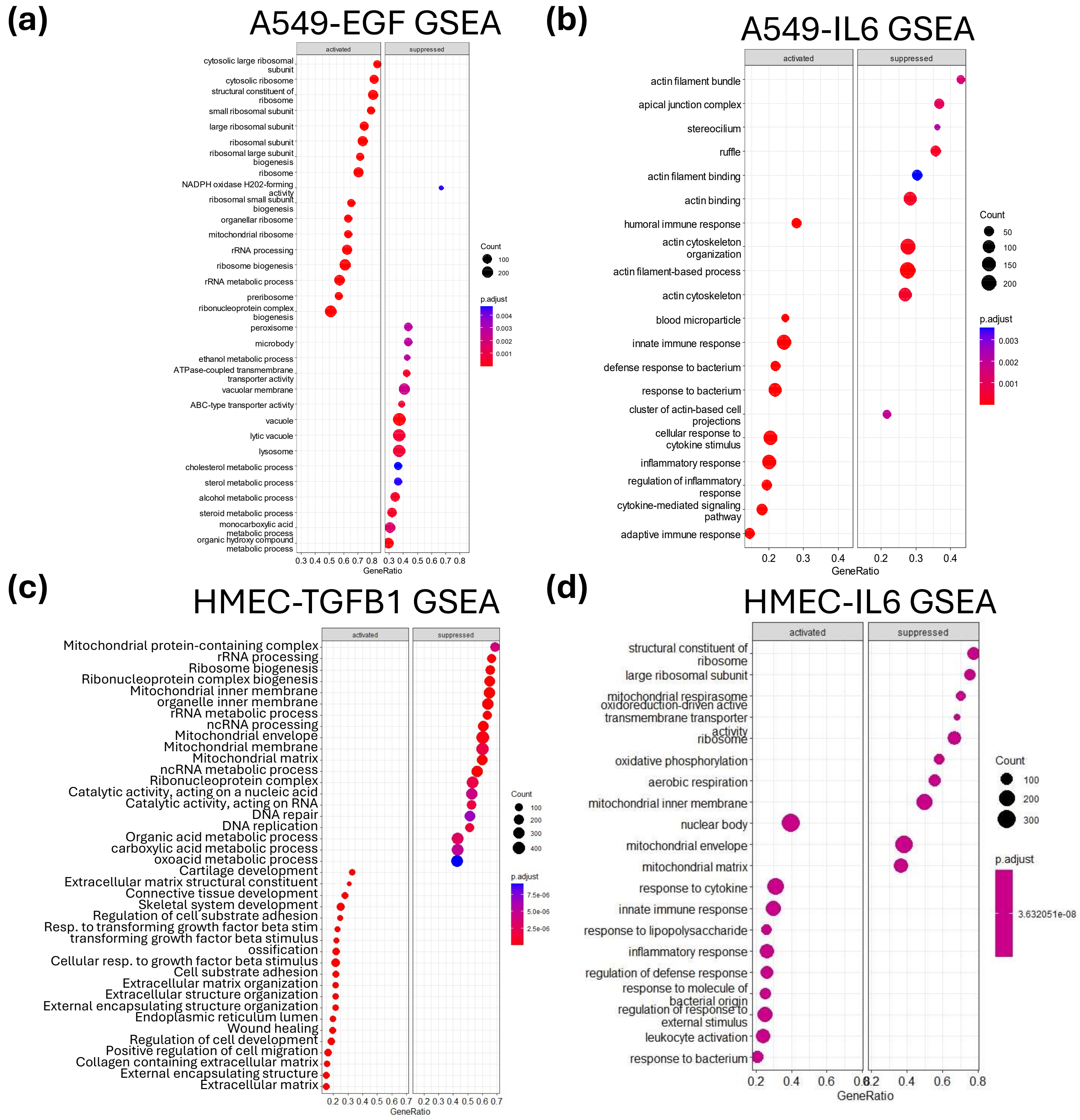


**Supplemental Figure 6.** GO GSEA analysis of responses to treatment in lung cancer and breast HMEC cells. (**A**) Dotplot of gene sets that were activated or repressed post treatment of A549 cells with EGF1. (**B**) Dotplot of gene sets that were activated or repressed post treatment of A549 cells with IL6. (**C**) Dotplot of gene sets that were activated or repressed post treatment with TGFB1 in the HMEC. (**D**) Dotplot of HMEC gene sets that were activated or repressed post treatment with IL-6 in the HMEC.
